# Integrating molecular markers and environmental covariates to interpret genotype by environment interaction in rice (*Oryza sativa L.*) grown in temperate areas

**DOI:** 10.1101/453712

**Authors:** Eliana Monteverde, Lucía Gutierrez, Pedro Blanco, Fernando Pérez de Vida, Juan E. Rosas, Victoria Bonnecarrère, Gastón Quero, Susan McCouch

## Abstract

Understanding the genetic and environmental basis of genotype × environment interaction (G×E) is of fundamental importance in plant breeding. If we consider G×E in the context of genotype × year interactions (G×Y), predicting which lines will have stable and superior performance across years is an important challenge for breeders. A better understanding of the factors that contribute to the overall grain yield and quality of rice (*Oryza sativa* L.) will lay the foundation for developing new breeding and selection strategies for combining high quality, with high yield. In this study, we used molecular marker data and environmental covariates (EC) simultaneously to predict rice yield, milling quality traits and plant height in untested environments (years), using both reaction norm models and partial least squares (PLS), in two rice breeding populations (*indica* and tropical *japonica*). We also sought to explain G×E by differential quantitative trait loci (QTL) expression in relation to EC. Our results showed that PLS models trained with both molecular markers and EC gave better prediction accuracies than reaction norm models when predicting future years. We also detected several milling quality QTL that showed a differential expression conditional on humidity and solar radiation, providing insight for the main environmental factors affecting milling quality in temperate rice growing areas.

## Introduction

Genetic by environment interaction (G×E) could be expressed as a difference in the relative response of genotypes across diverse environments. When we consider a set of genotypes exposed to different environments, their performance will differ depending on the interaction of genetic properties with the different environmental conditions, leading to differences in variances and rank changes among genotypes (Cooper and DeLacy 1994). These rank changes represent a very important challenge for breeders due to the difficulties of selecting genotypes with stable performance over diverse environments.

Environments can be different both in time and space. For this reason, the concept of G×E embraces both interactions that take place between genotypes and a particular location (genotype by location interaction), and between genotypes and particular years (genotype by year interaction). Genotype by location interactions are usually determined by soil and climate conditions and are thus predictable, while genotype by year interactions are characterized by plot-to-plot variability and weather conditions and thus, are harder to predict (Malosetti et al. 2013, 2016; van Eeuwijk et al. 2016). Several statistical approaches have been proposed to describe G×E in the context of classical plant breeding, including linear-bilinear models (Yates and Cochran 1938; Finlay and Wilkinson 1963; Gauch 1992; Crossa and Cornelius 1997) and mixed models with different covariance structures (Piepho 1998; Smith et al. 2001; Burgueño et al. 2007). Within linear models, factorial regression models are of particular importance because they permit the modeling of genotype sensitivity to specific environmental covariates (ECs) (van Eeuwijk et al. 1996; Vargas et al. 1998; Malosetti et al. 2004; Malvar et al. 2005). These models can also predict phenotypic responses in unobserved environments by using explicit environmental information.

Recent developments in sequencing technologies and statistical modeling have made it possible to use dense genotypic information to predict phenotypic responses through genomic prediction (GP). This idea was introduced by Meuwissen et al. (2001), and provides an alternative approach to indirect selection in crop breeding. GP models were originally developed for traits evaluated in single environments, but more recently standard GP models have been extended to account for G×E. Burgueño et al. (2012) were the first to extend genomic best linear unbiased prediction (GBLUP) to a multi-environment context, were G×E was modeled using genetic correlations. Lopez Cruz et al. (2015) proposed a GBLUP approach that explicitly models G×E interactions between all available markers and environments (M×E).

Standard GP models can be modified to accommodate climate information in the form of ECs. However, including ECs in the analysis can pose the same constraints encountered when using GP methods. As each covariate explains a small amount of the total environmental variance, a high number of ECs usually brings up multicollinearity issues. Several studies have proposed different ways to overcome multicollinearity, showing that the incorporation of explicit environmental and genetic information can improve model performance and predict performance in untested environments (Heslot et al. 2014; Jarquín et al. 2014; Malosetti et al. 2016). Jarquín et al. (2014) proposed a Bayesian reaction norm model where the main genetic and environmental effects are modeled using covariate structures that are functions of molecular markers and ECs respectively. The interaction effects between markers and ECs are modeled using a multiplicative operator. This model has the limitations that: 1) The Gaussian prior assumed for genetic and environmental effects does not induce variable selection, and 2) the model relies on the similarity between covariates through the relationship matrix, which may not be a good representation of reality. Heslot et al. (2014) proposed a factorial regression model, where instead of using all the available ECs and molecular markers, they chose the ECs that most significantly influenced the growth and development of the crop by using crop growth models (CGM). These variables were introduced in the factorial regression model along with those markers that showed the highest variance across environments. One advantage of this approach is that it reduces the dimensionality of both markers and EC; however, it is possible to lose information by using only preselected markers and ECs (Elias et al 2016).

The partial least square regression (PLS) (Wold et al. 2001) is a generalization of multiple linear regression (MLR). PLS is a dimension reduction approach that can accommodate a large number of correlated genetic and environmental variables simultaneously, by finding one or few factors named latent variables (LV) that explain both the variance of the between matrices **X** matrix (containing predictor variables) and the covariance **X** and **Y** (containing response variables). PLS can be used for variable selection, in order to improve estimation/prediction performance, but also to improve model interpretation and understanding of the system studied. PLS models have previously been tested in the context of GP both in plant and animal breeding (Solberg et al. 2009; Long et al. 2011; Colombani et al. 2012; Iwata et al. 2015), and in the context of G×E to detect highly influential environmental and marker covariates (Vargas et al. 1998; Crossa et al. 1999; Vargas et al. 1999).

Understanding the genetic basis of G×E is also necessary to gain predictive capability, and one way to do this is by detecting QTLs with varying effects across different environmental conditions, or QTL by environment interaction (QTL×E). Methods usually employed to detect QTL×E have been very useful to detect QTL with differential expression across environments, but provide no explanation of the underlying environmental factors involved. When weather data are available, factorial regression models can be used to determine the extent of influence of these factors on QTL×E (Crossa et al. 1999; Campbell et all. 2004; Malosetti et al. 2004).

Rice is one of the world’s most important staple food crops, constituting over 21% of the caloric intake of the world’s population and up to 76% of the caloric needs in many Asian countries (Fitzgerald et al. 2008). World markets dictate the value of rice mainly based on milling quality traits, so breeding for both high yield and quality is a major breeding objective for rice exporting countries like Uruguay. The main objectives of this study were to: 1) use molecular marker data and environmental covariates simultaneously to predict rice yield and milling quality traits in untested environments (years), and 2) Detect marker by environment covariate interactions that provide explanations of variable QTL effects across environments. Two rice breeding populations (*indica* and *tropical japonica*) were used in this study and were evaluated for grain yield, plant height and grain quality traits (head rice percentage and chalky grain percentage) across 3-5 years in Eastern Uruguay. Results from these two analyses provided clues about the main environmental variables that could be driving G×E in temperate rice-growing regions such as Uruguay.

## Materials and Methods

### Germplasm

The germplasm consists of two rice-breeding populations, an *indica* and a *tropical japonica* population belonging to the National Institute of Agricultural Research (INIA-Uruguay). Both populations were evaluated in a single location, Paso de la Laguna Experimental Station (UEPL), Treinta y Tres, Uruguay (33°15’S, 54°25’W) over between 2009-2013.

The *indica* population consisted of 327 elite breeding lines evaluated over three years (2010-2012), and the field design consisted in a randomized complete block design with two or three replications. Trait correlations, heritabilities, and genomic prediction accuracies for this dataset were computed in previous studies (Rosas et al. 2017, Monteverde et al. 2018, Quero et al. 2018). The *tropical japonica* population consisted of 320 elite breeding lines evaluated over five years (2009-2013). The number of accessions observed each year ranged from 93 to 319, as detailed in Table 1. This dataset was unbalanced with non-random missing data, since ~50% genotypes were dropped from testing every year based on performance, and new genotypes were added over time. Each year, the genotypes were planted independently in replicated trials in six-row plots using an augmented randomized complete block design with two or three replications. Both *indica* and *tropical japonica* trials were conducted under irrigated conditions using appropriate pest and weed control.

**Table 1:**
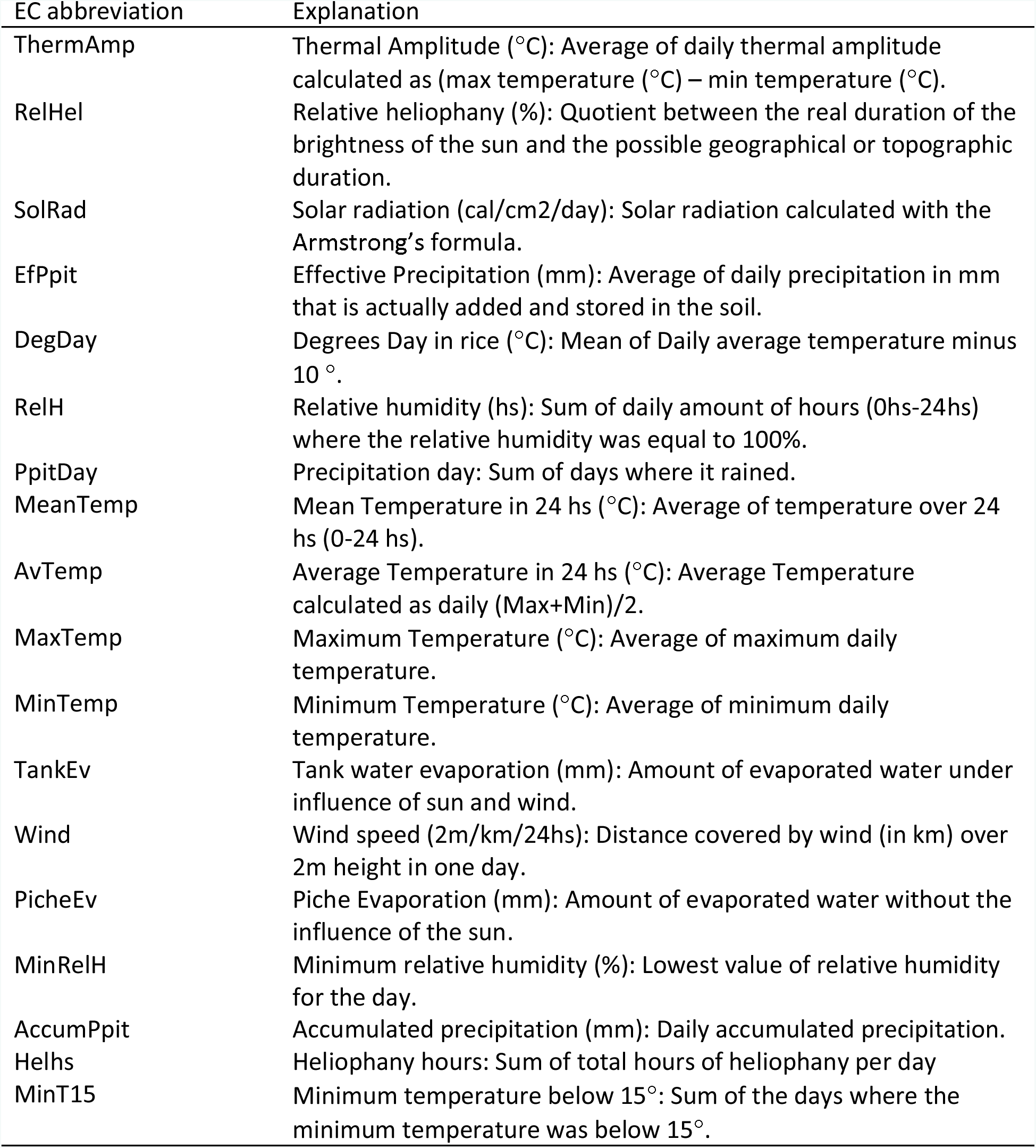
Environmental covariates used in this study.

| EC abbreviation | Explanation |
| --- | --- |
| ThermAmp | Thermal Amplitude (°C): Average of daily thermal amplitude calculated as (max temperature (°C) – min temperature (°C)). |
| RelHel | Relative heliophany (%): Quotient between the real duration of the brightness of the sun and the possible geographical or topographic duration. |
| SolRad | Solar radiation (cal/cm<sup>2</sup>/day): Solar radiation calculated with the Armstrong's formula. |
| EfPpit | Effective Precipitation (mm): Average of daily precipitation in mm that is actually added and stored in the soil. |
| DegDay | Degrees Day in rice (°C): Mean of Daily average temperature minus 10 °. |
| RelH | Relative humidity (hs): Sum of daily amount of hours (0hs-24hs) where the relative humidity was equal to 100%. |
| PpitDay | Precipitation day: Sum of days where it rained. |
| MeanTemp | Mean Temperature in 24 hs (°C): Average of temperature over 24 hs (0-24 hs). |
| AvTemp | Average Temperature in 24 hs (°C): Average Temperature calculated as daily (Max+Min)/2. |
| MaxTemp | Maximum Temperature (°C): Average of maximum daily temperature. |
| MinTemp | Minimum Temperature (°C): Average of minimum daily temperature. |
| TankEv | Tank water evaporation (mm): Amount of evaporated water under influence of sun and wind. |
| Wind | Wind speed (2m/km/24hs): Distance covered by wind (in km) over 2m height in one day. |
| PicheEv | Piche Evaporation (mm): Amount of evaporated water without the influence of the sun. |
| MinRelH | Minimum relative humidity (%): Lowest value of relative humidity for the day. |
| AccumPpit | Accumulated precipitation (mm): Daily accumulated precipitation. |
| Helhs | Heliophany hours: Sum of total hours of heliophany per day |
| MinT15 | Minimum temperature below 15°: Sum of the days where the minimum temperature was below 15°. |

The agronomic traits of interest used in this study were Grain Yield (GY of paddy rice in kilograms per hectare) and Plant Height (PH measured in cm from the soil surface to the tip of the flag leaf). The grain quality traits measured were Percentage of Head Rice Recovery (PHR measured in grams, as the weight of whole milled kernels, using a 100g sample of rough rice), and the percentage of Chalky Grain (GC measured as % of chalky kernels in a subsample of 50 g of total milled rice, where the area of chalk - core, white back or white belly - was larger than half the kernel area based on visual inspection). More details about how grain quality traits were measured can be found in Quero et al. (2017) and Monteverde et al. (2018).

### Phenotypic analysis

Phenotypic data for each trait were analyzed separately each year. The model used to calculate the best linear unbiased estimators (BLUEs) for each year was:

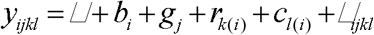

where *y*_*ijkl*_ is the trait score, ▱ is the overall mean, *b*_*i*_ is the random block effect with 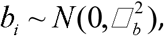 where 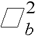 is the block variance, *g*_*j*_ is the genotypic effect, *r*_*k*(*i*)_ and *c*_*l*(*i*)_ are the random row and column effects nested within blocks with 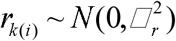 and 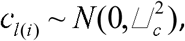 where 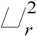 and 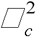 are the row and column variances respectively, and ▱_*ijkl*_ is the model residual vector with 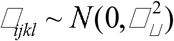 where 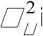 is the error variance.

### Genotypic characterization

The lines were genotyped using genotyping-by-sequencing (GBS). SNP calling was performed using the TASSEL 3.0 GBS pipeline (Bradbury et al. 2017), and SNPs were aligned to the Nipponbare reference genome MSU version 7.0 (http://rice.plantbiology.msu.edu/) using Bowtie 2 (Langmead and Salzberg, 2012). Imputation of missing data was performed with the FILLIN algorithm implemented in TASSEL 5.0 (Swarts et al. 2008) for both datasets separately. The GBS datasets were filtered to retain markers with <50% missing data after imputation, and a minor allele frequency MAF<0.05, as reported by Quero et al. (2018), and Monteverde et al. (2018). The final *indica* and *tropical japonica* marker dataset consisted of 92,430 and 44,598 SNP markers respectively.

### Derivation of EC from weather data

Daily weather data were obtained from GRAS unit from INIA (http://www.inia.uy/gras/Clima/Banco-datos-agroclimatico). The database contains weather data from 1965 to the last calendar month completed, for all 5 INIA experimental stations in Uruguay. The variables available were related to temperature, precipitation, solar radiation, humidity, wind, and evaporation.

To compute the EC from daily weather data for each rice genotype, the plant development stage has to be determined in order to account for the differential effect that weather variables may have in different stages of crop development. This information is usually hard to obtain directly or, as in our case, not available. For this reason, the phenology of the crop was defined according to flowering time (days to 50% flowering, FT), which was measured for each line every year. With this measure, and sowing and harvest date, three main phenology stages were determined for each year, according to Yoshida (1981): vegetative stage (of variable length, starting on sowing date), reproductive stage (starting 35 days before FT), and maturation stage (ending 30 days after FT).

Once these phenological stages are defined for each year, EC can be computed from daily weather data. Covariates with zero variance were removed from the analysis. For prediction, both markers and EC were centered by subtracting the mean, and standardized to unit variance by dividing the centered values by the standard deviation of the marker or EC. A total of 54 EC were used in the *indica* population, and 51 for the tropical *japonica*, and are summarized in Table 2.

**Table 2:** Description of the rice breeding lines evaluated each year and broad-sense heritabilities for each trait calculated in a line-basis

| *indica* |  |  |  |  |  |
| --- | --- | --- | --- | --- | --- |
| Year | Lines evaluated | $H^2$ | | | |
|  |  | GY | PHR | GC | PH |
| 2010 | 327 | 0.46<sup>a</sup> | 0.86<sup>a</sup> | 0.73<sup>a</sup> | 0.49<sup>a</sup> |
| 2011 | 327 | 0.60<sup>a</sup> | 0.78<sup>a</sup> | 0.59<sup>a</sup> | 0.66<sup>a</sup> |
| 2012 | 327 | 0.68<sup>a</sup> | 0.71<sup>a</sup> | 0.69<sup>a</sup> | 0.58<sup>a</sup> |
| *Tropical japonica* |  |  |  |  |  |
| 2009 | 93 | 0.44 | 0.67 | 0.59 | 0.77 |
| 2010 | 292 | 0.68 | 0.71 | 0.59 | 0.62 |
| 2011 | 319 | 0.43<sup>a</sup> | 0.85<sup>a</sup> | 0.41<sup>a</sup> | 0.62<sup>a</sup> |
| 2012 | 319 | 0.57<sup>a</sup> | 0.79<sup>a</sup> | 0.75<sup>a</sup> | 0.79<sup>a</sup> |
| 2013 | 134 | 0.70 | 0.75 | 0.80 | 0.78 |

<sup>1</sup> Previously reported by Monteverde et al. (2018) and Rosas et al. (2017)

### PLS regression

PLS regression was first introduced by Wold (1966), and was originally developed for econometrics and chemometrics. It is a multivariate statistical technique that was designed to deal with the *p* > > *n* problem; i.e., when the number of) is much larger (and more highly correlated) than the number of observations (explanatory variables (*p*). A brief explanation of PLS relating one response variable (*y*) to a set of explanatory variables (*X*) is given below, but it can be extended to more than one response variable (Boulesteix and Strimmer 2006; Wold 2001).

In PLS, the data for *p* explanatory variables are given by the matrix **X** = (**x_1_**,…, **x**_*p*_), and data for the dependent variables are given by the response vector **y**. Each x_1_, …, x_p_ and **y** vectors have *n* × 1 dimensions corresponding to the number of observations. In this work, the **y** vector contains all the observations for a given trait in different environments (years), and the columns of the **X** matrix are the variables corresponding to either markers only, or markers and ECs. All variables in PLS must be centered and scaled.

PLS is based on the latent variable (LV) decomposition:

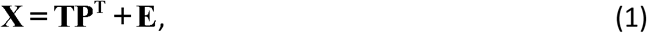

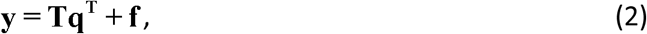

where **T** is a *n* × *c* matrix giving the LV (also called scores) for the *n* observations, and **P** (*p* × *c*) is a matrix of *p*-dimensional orthogonal vectors called *X*-loadings, **q** (1 × *c*) is a vector of scalars and, also named *Y*-loadings, **E** (*n* × *p*) and **f** (*n* × 1) are a residual matrix and vector respectively.

The LV matrix **T** that relates the **X** matrix to the vector **y** is calculated as:

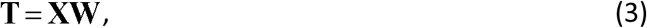

where **W**, is a (*p* × *c*) matrix of weights. For a given matrix **W**, the LV obtained by forming corresponding linear transformations of the variables in **X**, *X*_1_, …, *X*_*p*_ are denoted as *T*_1_, …, *T*_*c*_:

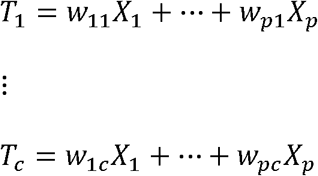

These LV are then used for prediction in place of the original variables. After computing the **T** matrix, **q^T^** is obtained as the least squares solution of Eq. (2):

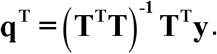

The vector **b** of regression coefficients for the model **y = Xb+ f**, to predict new responses, is calculated as:

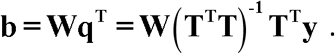

Since regression and dimension reduction are performed simultaneously, all **b**, **T**, **W**, **P** and **q** are part of the output. Both **X** and **y** are taken into account when calculating the LV in **T**. Moreover, they are defined so that the covariance between the LV and the response is maximized.

In PLS, the optimal number of LV (*c*) must be determined. In this work, we used the root means squared error of prediction (RMSEP),

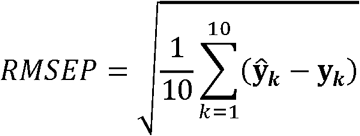

which was minimized with 10-fold cross-validation in the training data set and for each value of LV (Mevik and Cederkvist, 2004). PLS models calculations were performed with the R package “mixOmics” (Lê Cao et al. 2016).

### Genomic Best Linear Prediction (GBLUP) and reaction norm models

Mixed linear models were used as a baseline comparison of prediction accuracies with PLS models. The models used considered the random main effects of markers (G model), the random main effects of markers and EC (G+W model), and the random main effects of markers, EC, and the interactions between them (G+W+GW model).

The G model constituted of a standard GBLUP model for the mean performance of genotypes within each set of environments, using the following model:

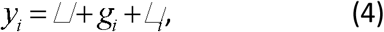

where ▱ is the overall mean, *g*_*i*_ is the genotypic random effect of the *i*^*th*^ line expressed as a regression on marker covariates of the form: 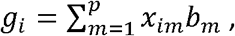 where *x*_*im*_ is the genotype of the *i*^*th*^ line at the *m*^*th*^ marker, and *b*_*m*_ is the effect of the *m*^*th*^ marker. Marker effects are considered as IID draws from normal distributions of the form 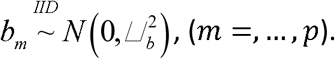

The vector **g = Xb** contains the genomic values of all the lines, and follows a multivariate normal density with null mean and covariance matrix 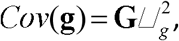 where **G** is a genomic relationship matrix whose entries are given by **G**= **XX^T^**/*p*.

As previously reported by Jarquín et al. (2014), it is possible to model the environmental effects with a random regression on the EC that describes the environmental conditions faced by each genotype, that is: 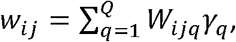 where *W*_*ijq*_ is the value of the *q*^*th*^ EC evaluated in the *ij*^*th*^ environment × genotype combination, ▱_*q*_ is the main effect of the corresponding EC, and *Q* is the total number of EC. Again, we consider the effects of the EC as IID draws from normal densities, 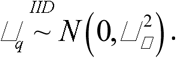 The vector **w**= **W**▱ follows a multivariate normal density with null mean and a covariance matrix proportional to Ω whose entries are computed the same way as those of the **G** matrix but using EC instead of markers. This covariance structure describes the similarity among environmental conditions. Then, the model becomes:

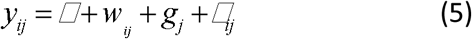

This model also includes a marker × EC interaction term, where the covariance of the interaction is modeled by the Hadamard product of 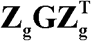 and **Ω**, denoted as 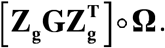. This model extends Eq. (4) as follows:

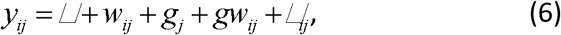

with 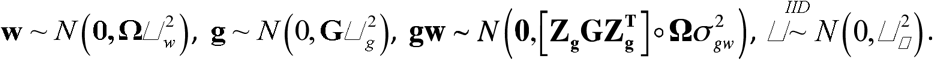

### Assessing prediction accuracy for new environments

The prediction problem studied here was that of predicting future seasons. This prediction was performed by using one year in the testing population, and the rest of the years in the training population. Prediction accuracies obtained from both PLS and Reaction norm models were assessed by calculating the Pearson correlation between the predicted values from each model, and the observed values.

### QTL by EC interactions

For the detection of QTL by environment interaction we used a two-step strategy as described in Gutiérrez et al. (2015). In the first step, we scanned the genome of both *indica* and *tropical japonica* subspecies to detect QTL in individual environments (single environment QTL mapping). In the second step, QTL expression across environments was regressed on environmental covariates in order to explain QTL effects in terms of sensitivities to environmental covariates (Malosetti et al. 2004; Boer et al. 2007; Malosetti et al. 2013).

For the first step, we fitted a mixed model for single environment QTL detection. The model used was the kinship model with:

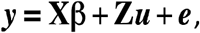

where ***y*** is the vector of phenotypic means for that environment, **X** is the molecular marker score matrix, **β** is the vector of marker effects, **Z** is an incidence matrix, ***u*** is the vector of random background polygenic effects with variance 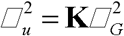 (where **K** is the kinship matrix, and 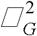 is the genetic variance), and ***e*** is the vector of residuals. A GWAS analysis for each dataset, trait and environment was performed using the R statistical software (R Core Team, 2017) with the package GWASpoly (Rosyara et al. 2016) fitting the additive model. For QTL determination in each environment, we used the Benjamini-Hochberg FDR (α=0.05) to control the type I error (Benjamini and Hochberg 1995).

In the second step, all marker-trait associations detected in the first step were fitted in a second mixed model assuming a linear relationship between the effect of the QTL and a given environmental covariate, using the model presented in Malosetti et al. (2013) given by:

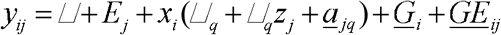

where *y*_*ij*_ is the phenotype of individual *i* at environment *j*, ▱ is the general mean, *E_j_* is the environmental main effect, *x*_*i*_ is the value of the marker predictor, α_*q*_ is the effect of QTL *q* in the average environment, β_*q*_ corresponds to the change of the QTL effect per unit of change of the covariable’s value, and *<u>a</u>*_*iq*_ is the random effect corresponding to the residual (unexplained) QTL effect, with 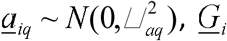 is the random remaining (not due to the QTL) genotype effect with 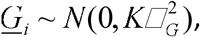 and *<u>GE</u>*_*ij*_ is the remaining (random) G×E effect, with 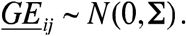 Three different models for the variance-covariance matrix ▱ were compared: compound symmetry (CS) where the genetic variances are homogeneous across environments 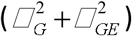 and the genetic covariances between environments are modeled by 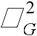; heterogeneous compound symmetry (HCS), which allows for heterogeneous genetic variances across environments 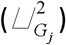 and a common genetic covariance parameter 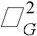; and the unstructured (UN) model with a specific genetic variance parameter per environment and a specific genetic covariance between environment. The different models were compared using the Bayesian information criterion (BIC) to select the optimal model (Broman and Speed 2002). We tested for the significance of the fixed terms in mixed models using Wald test at a *p* value of 0.05, following Malosetti et al. (2004). Mixed models for QTL×EC interaction were computed with the R package *sommer* (Covarrubias 2016).

### Data Availability

All the data used in this study are provided in Supplemental Material. Genotype data can be found as RDS files (“geno_indica.rds” and “geno_japonica.rds”), phenotypic data can be found as “.csv “ files (“pheno_indica.csv” and “pheno_japonica”), marker positions can be found in files “map_indica.csv” and “map_japonica.csv”, and EC data are available in files “EC_indica.csv” and “EC_japonica.csv”.

## Results

### Phenotypic data analysis

The *indica* dataset was balanced with a total of 327 lines per environment, while the *tropical japonica* dataset was unbalanced, with a total of 23 lines common to all environments (Table 1). Estimations of broad-sense heritability estimated on a line-mean basis per trait by year for both datasets were medium to high, with PHR having the highest values of heritability in both datasets.

Table 3 shows the partitioning of the observed phenotypic variance into different sources of variation for both rice datasets. In the *indica* population, PHR and GC showed the highest proportion of variance explained by G×Y, at 20.04% and 13.22%, respectively. On the other hand, the year component was the highest variance component for GY and PH (Table 3). In the *tropical japonica* population, the year component was the highest; it was above all components for the four traits, and much higher than for the *indica* population. In contrast, the G×Y component was lower in *tropical japonica* compared to *indica* (Table 3).

**Table 3:** Trait variance component estimation and proportion of the total variance explained for the four traits evaluated in Uruguayan *indica* and *tropical japonica* populations.

| *indica* |  |  |  |  |  |
| --- | --- | --- | --- | --- | --- |
| GY |  |  | PHR |  |  |
| Group | Variance | % | Group | Variance | % |
| Year | 496540 | 18.9 | Year | 0.0001 | 10.02 |
| Genotype | 379554 | 14.5 | Genotype | 0.0002 | 20.04 |
| GxY | 143374 | 5.5 | GxY | 0.0002 | 20.04 |
| Column | 55361 | 2.1 | Column | 0.000007 | 0.70 |
| Row | 31357 | 1.2 | Row | 0.000006 | 0.60 |
| Block | 516792 | 19.7 | Block | 0.0002 | 20.04 |
| Residual | 1000130 | 38.1 | Residual | 0.000285 | 28.56 |
| GC |  |  | PH |  |  |
| Group | Variance | % | Group | Variance | % |
| Year | 0.0003 | 13.22 | Year | 12.51 | 9.26 |
| Genotype | 0.0004 | 17.62 | Genotype | 8.91 | 6.60 |
| GxY | 0.0003 | 13.22 | GxY | 5.56 | 4.12 |
| Column | 0.0003 | 13.22 | Column | 0.84 | 0.62 |
| Row | 0.00007 | 3.08 | Row | 0.92 | 0.68 |
| Block | 0.0005 | 22.03 | Block | 8.31 | 6.15 |
| Residual | 0.0004 | 17.62 | Residual | 98.02 | 72.57 |
| *tropical japonica* |  |  |  |  |  |
| GY |  |  | PHR |  |  |
| Group | Variance | % | Group | Variance | % |
| Year | 1988252 | 43.2 | Year | 0.001 | 41.36 |
| Genotype | 197961 | 13.2 | Genotype | 0.0003 | 12.41 |
| GxY | 99052 | 5.1 | GxY | 0.0001 | 4.14 |
| Column | 24932 | 0.3 | Column | 0.000008 | 0.33 |
| Row | 19062 | 0.5 | Row | 0.00001 | 0.41 |
| Block | 115775 | 22.7 | Block | 0.0006 | 24.81 |
| Residual | 682613 | 15.1 | Residual | 0.0004 | 16.54 |
| GC |  |  | PH |  |  |
| Group | Variance | % | Group | Variance | % |
| Year | 0.005 | 66.32 | Year | 37.5 | 43.7 |
| Genotype | 0.0007 | 9.29 | Genotype | 18.4 | 21.4 |
| GxY | 0.0006 | 7.96 | GxY | 1.6 | 1.8 |
| Column | 0.000009 | 0.12 | Column | 0.1 | 0.1 |
| Row | 0.00003 | 0.40 | Row | 0.1 | 0.2 |
| Block | 0.0004 | 5.31 | Block | 13.3 | 15.5 |
| Residual | 0.0008 | 10.61 | Residual | 14.8 | 17.2 |

### Genomic prediction of untested years

Bar plots showing prediction accuracy for the four traits in the *indica* population are shown in Figure 1. PLS-based methods showed higher prediction accuracies than reaction norm-based models for all traits except GC, where prediction accuracies for the PLS-GW method were the same as with the reaction norm models. For PLS models, the use of EC in addition to molecular markers resulted in higher prediction accuracies in all cases, though PHR in 2011 and GC in 2012 had identical prediction accuracies for both methods. For reaction norm models, fitting the G+W+GW model resulted in either lower or equal prediction accuracies than fitting the simpler G+W model (Fig. 1).

**Figure 1:**
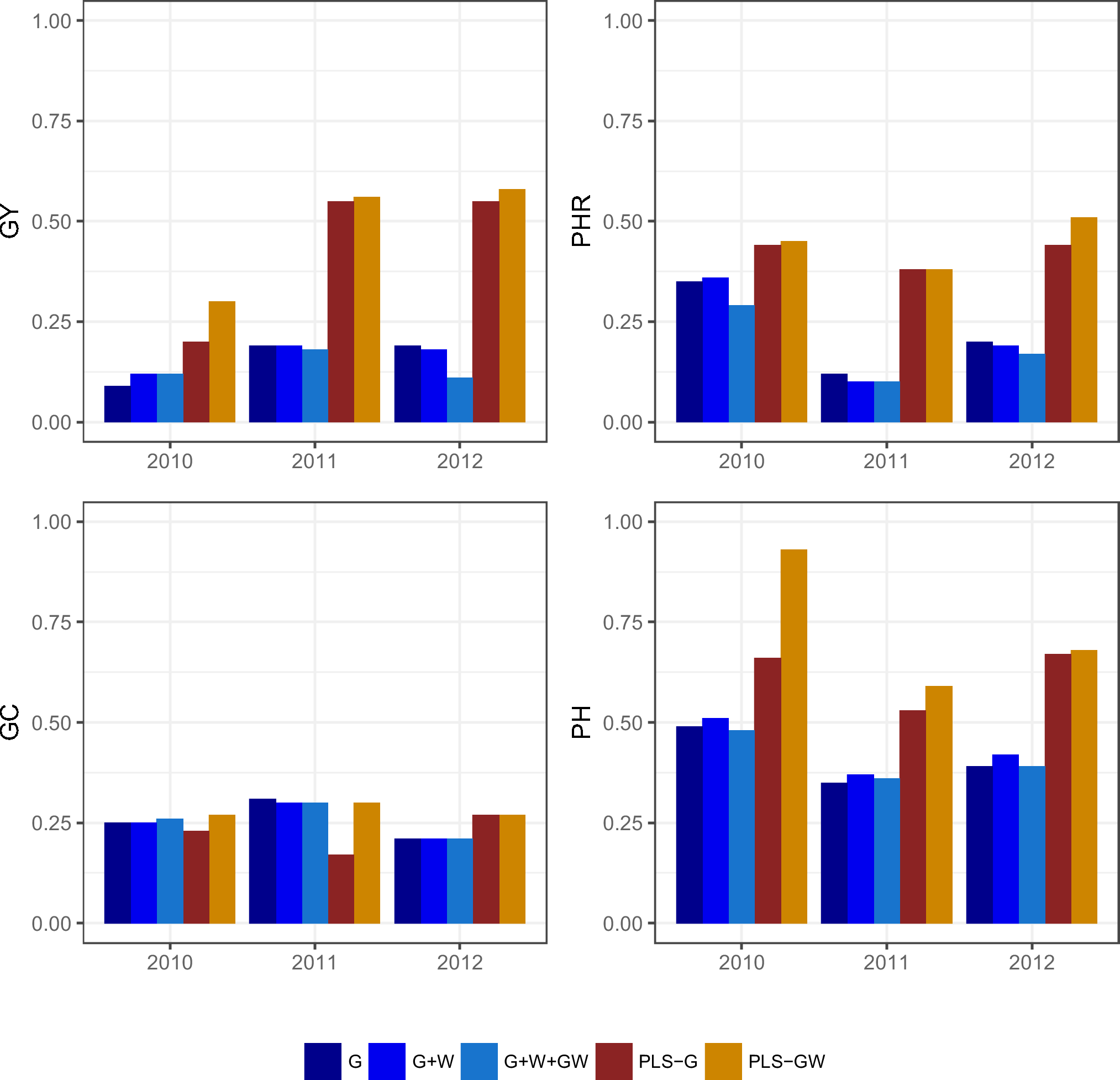
Correlations between predicted vs. observed values for Grain Yield (GY), Head Rice Percentage (PHR), Grain Chalkiness percentage (GC) and Plant Height (PH) for new years with the G, G+W, G+W+GW, PLS-G and PLS-GW for the *indica* rice breeding population.

In the *tropical japonica* population, the use of PLS-based models was always better than reaction norm models, with the single exception of GY in 2010 (Fig. 2). In all cases, PLS-GW was better than PLS-G. Within the reaction norm models, the G+W method was the best, with the exception of GC in 2013. Fitting a G×E component in these models resulted in lower prediction accuracies than fitting the G+W model (Fig. 2).

**Figure 2:**
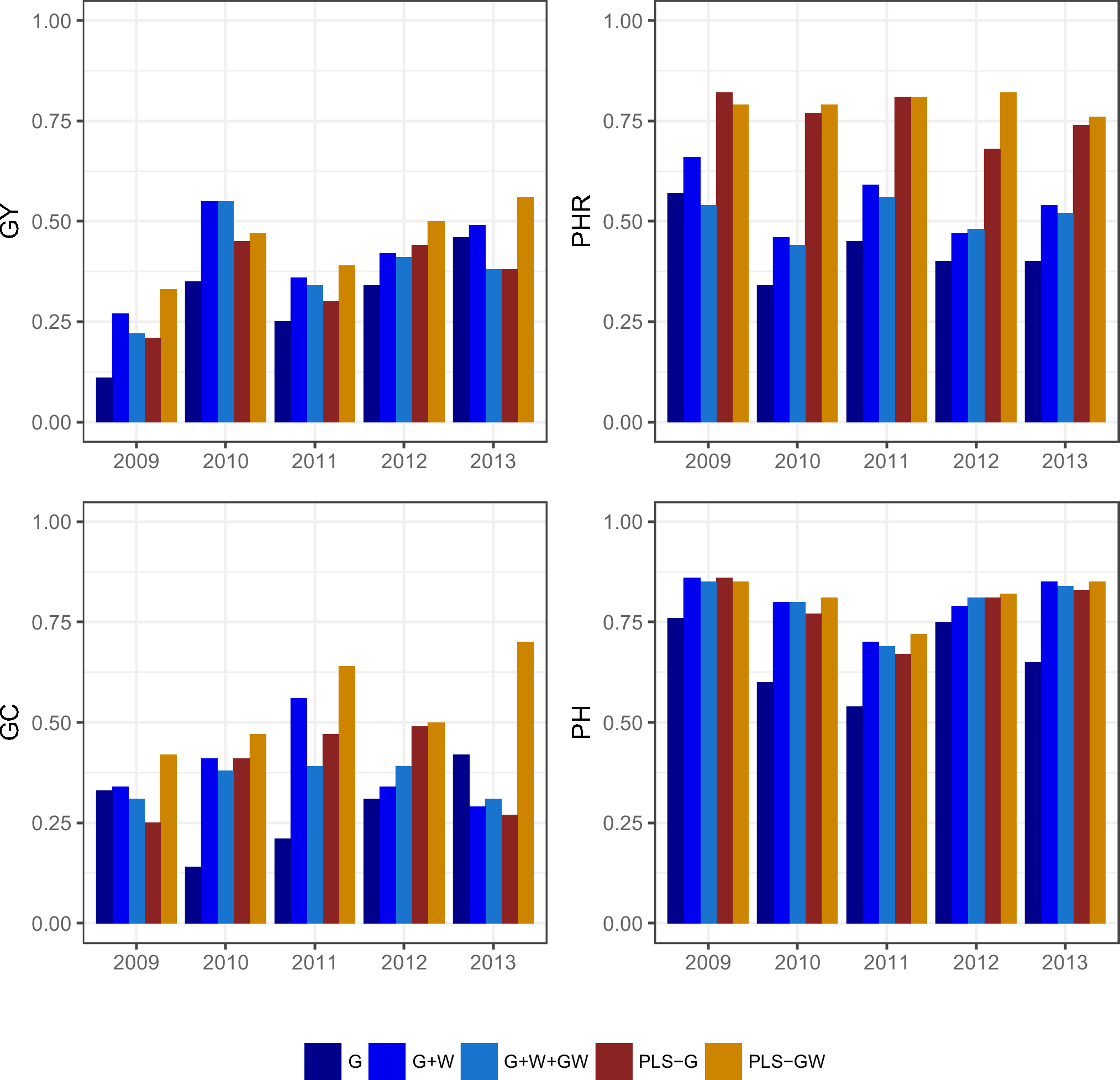
Correlations between predicted vs. observed values for Grain Yield (GY), Head Rice Percentage (PHR), Grain Chalkiness percentage (GC) and Plant Height (PH) for new years with the G, G+W, G+W+GW, PLS-G and PLS-GW for the tropical *japonica* rice breeding population.

When running PLS with all the environments within each dataset, we can detect which variables best explain each trait by looking at the coefficients. Table 4 shows the ranking of coefficients for the EC variables for each trait in both datasets. For GY, variables related to temperature and humidity during flowering stage were among the most important. For PHR and GC, the 5 variables with the highest coefficients were related to temperature, humidity and solar radiation during maturation. In the *tropical japonica* dataset, variables related to humidity, solar radiation and rainfall during maturation showed the highest coefficients for PHR and GC. For GY, two variables at flowering time showed higher coefficient values than the rest: maximum temperature and wind speed (Table 4).

**Table 4:** Top 5 PLS-GW coefficients for the environmental covariates for Grain Yield (GY), Head Rice Percentage (PHR), Grain Chalkiness percentage (GC) and Plant Height (PH) for the *indica* and *tropical japonica* rice breeding populations.

| *indica* |  |  |  |  |  |
| --- | --- | --- | --- | --- | --- |
| Type | Variable | Coefficient |  |  |  |
|  |  | GY | PHR | GC | PH |
| Temperature | ThermAmp_V | - | 0.0001251 | -0.000403 | - |
|  | MinTemp_R | 0.005277 | - | - | -0.00751 |
|  | MeanTemp_R | 0.005265 | - | - | -0.00773 |
|  | ThermAmp_M | - | 0.0001273 | -0.000402 | - |
| Precipitation | EfPpit_R | 0.005159 | - | - | -0.00723 |
| Evaporation | TankEv_V | 0.005185 | - | - | -0.00772 |
|  | TankEv_M | 0.0001272 | - | -0.000407 | - |
| Humidity | MinRelH_M | - | -0.0001270 | 0.000406 | - |
| Radiation | SolRad_M | - | 0.0001267 | -0.000402 | - |
| Wind | Wind_V | -0.005108 | - | - | 0.00766 |
| *tropical japonica* |  |  |  |  |  |
| Type | Variable | Coefficient |  |  |  |
|  |  | GY | PHR | GC | PH |
| Temperature | MinTemp_V | - | - | - | -0.014 |
|  | MaxTemp_V | 0.0102 | - | - | -0.015 |
|  | MeanTemp_V | - | - | - | -0.016 |
|  | DegDayRice_V | - | - | - | -0.016 |
|  | MaxTemp_R | -0.0125 | - | - | - |
|  | AvTemp_M | -0.0101 | - | - | - |
| Precipitation | PpitDay_R | - | - | - | 0.014 |
|  | EfPpit_M | - | 0.013 | 0.018 | - |
|  | AccumPpit_M | - | -0.013 | - | - |
| Evaporation | PicheEv_V | - | 0.013 | - | - |
| Humidity | MinRelH_M | - | -0.015 | 0.019 | - |
| Radiation | Helhs_M | - | 0.013 | -0.018 | - |
|  | SolRad_M | - | - | -0.017 | - |
|  | RelHel_M | - | - | -0.017 | - |
| Wind | Wind_V | -0.0103 | - | - | - |
|  | Wind_R | -0.0126 | - | - | - |

### Detecting QTL in single environments

We searched for significant trait-marker associations in single years to find QTL to test for interactions with EC in the next step. In this first analysis, we could not find any QTL that passed the FDR threshold for GY in any environment in any population. In the case of PH, we did not find any QTL for the *indica* population, but we found one major effect QTL on chromosome 1 that was significant in all environments in the *japonica* dataset; it corresponds to the *sd-1* gene.

We detected QTL for grain quality traits in both datasets (Table 5). In the *indica* population, a total of 13 QTL (chromosomes 1, 2, 3, 4, 6, 7, 10 and 11) were found for PHR, and a total of 4 QTL (chromosomes 1, 3 and 4) for GC. QTL were found only in years 2010 and 2012 for PHR, and in years 2011 and 2012 for GC. Three of the QTL were reported in a previous GWAS analysis using this same dataset (Quero et al. 2018). These QTL were: qPHR.i.2.2 (S2_24210614), qPHR.i.3.1 (S3_10247958), qGC.i.1.1 (S1_1066894). Two additional QTL were in LD with two previously reported QTL in the same study. These were qPHR.i.3.2 and qPHR.i.6.1, which were in LD with S3_15365726 and S6_829223 in our study, respectively. In the *tropical japonica* population, a total of 5 QTL were found for PHR (chromosomes 1, 2, 3, 6, and 8), and one for GC (chromosome 6) (Table 5). Two of these QTL were in LD with previously reported QTL: qPHR.j.3.1 with S3_1395165, and qGC.j.6.2 with S6_27402260 (Quero et al. 2018). No significant QTL were found for GY or PH for any year in either of the populations.

**Table 5:**
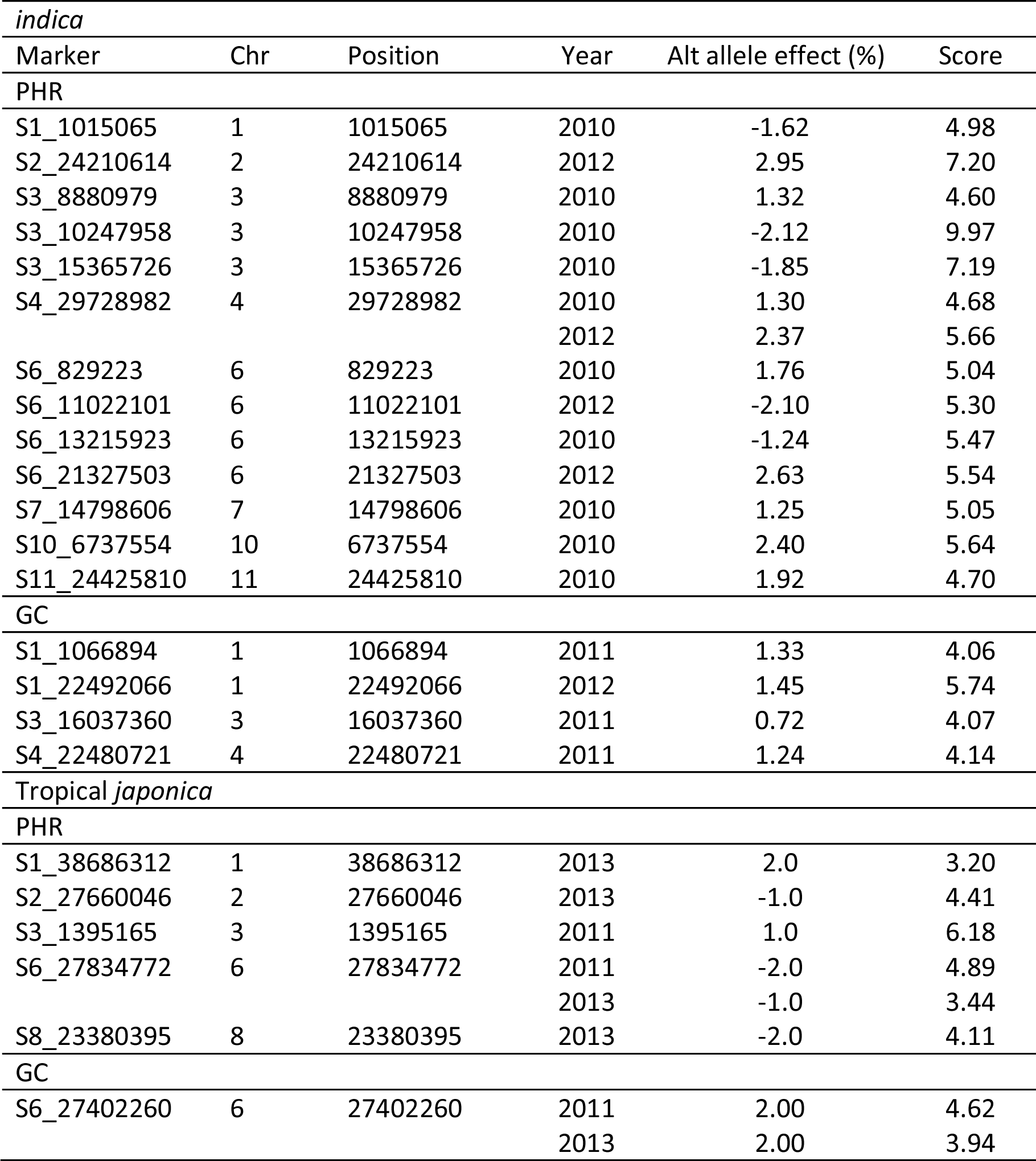
Marker-trait associations for PHR and GC traits in *indica* and *tropical japonica* rice breeding populations. Chromosome position (bp), year, effect of the alternative allele, and score (-log_10_(p-value)) are shown in the table.

### QTL × environmental covariate interactions

A decomposition of the QTL with significant QTL × environment interaction was obtained by introducing environmental covariates as explanatory variables. We first tested different covariance structures for the modeling of the G×E component and compared them using BIC (see Methods); for all traits, the covariance structure that resulted in the lowest BIC was the heterogeneous compound symmetry (data not shown). The QTL responses for the *indica* dataset are shown in Table 6. Two QTL showed significant interaction with environmental covariates related to temperature, precipitation and humidity during the maturation stage. Marker S2_24210614 showed a negative relationship with PpitDay_M and RelH_M. The high correlation between these two variables (*r* = 0.99) explains why they show the same coefficients for the main QTL effect (α), and the interaction (β). Marker S6_13215923 showed a negative relationship when interacting with low temperature days, and a positive effect when interacting with maximum and average temperature during maturity. There were not significant main effects detected at either of these positions (Table 6).

**Table 6:** QTL responses to EC for PHR in the *indica* rice population.

| Trait | Marker | Chromosome | Position | EC | $\alpha$ | $\beta$ |
| --- | --- | --- | --- | --- | --- | --- |
| PHR | S2_24210614 | 2 | 24210614 | PpitDay_M | 0.09 | -0.03* |
|  |  |  |  | RelH_M | 0.08 | -0.04* |
|  |  |  |  | MinTemp_R | 0.10 | -0.04* |
| PHR | S6_13215923 | 6 | 13215923 | Temp15_M | -0.09 | -0.05* |
|  |  |  |  | MaxTemp_M | -0.1 | 0.17* |
|  |  |  |  | AvTemp_M | -0.09 | 0.19* |

$\alpha$ : QTL main effect
$\beta$ : Slope parameter for the QTL×EC parameter
\* significance level at $\alpha = 0.05$

Results for regression of marker covariates on environmental covariates for the *tropical japonica* dataset are shown in Table 7. One QTL for PHR, S6_27834772, showed significant interaction with environmental covariates related to humidity and precipitation during maturation. This marker did not show a significant main effect (Table 7). For GC, marker S6_27402260, also in chromosome 6, showed a significant positive response to weather covariates related to precipitation and temperature, and a negative response to heliophany. This marker also showed a significant main effect (Table 7).

**Table 7:**
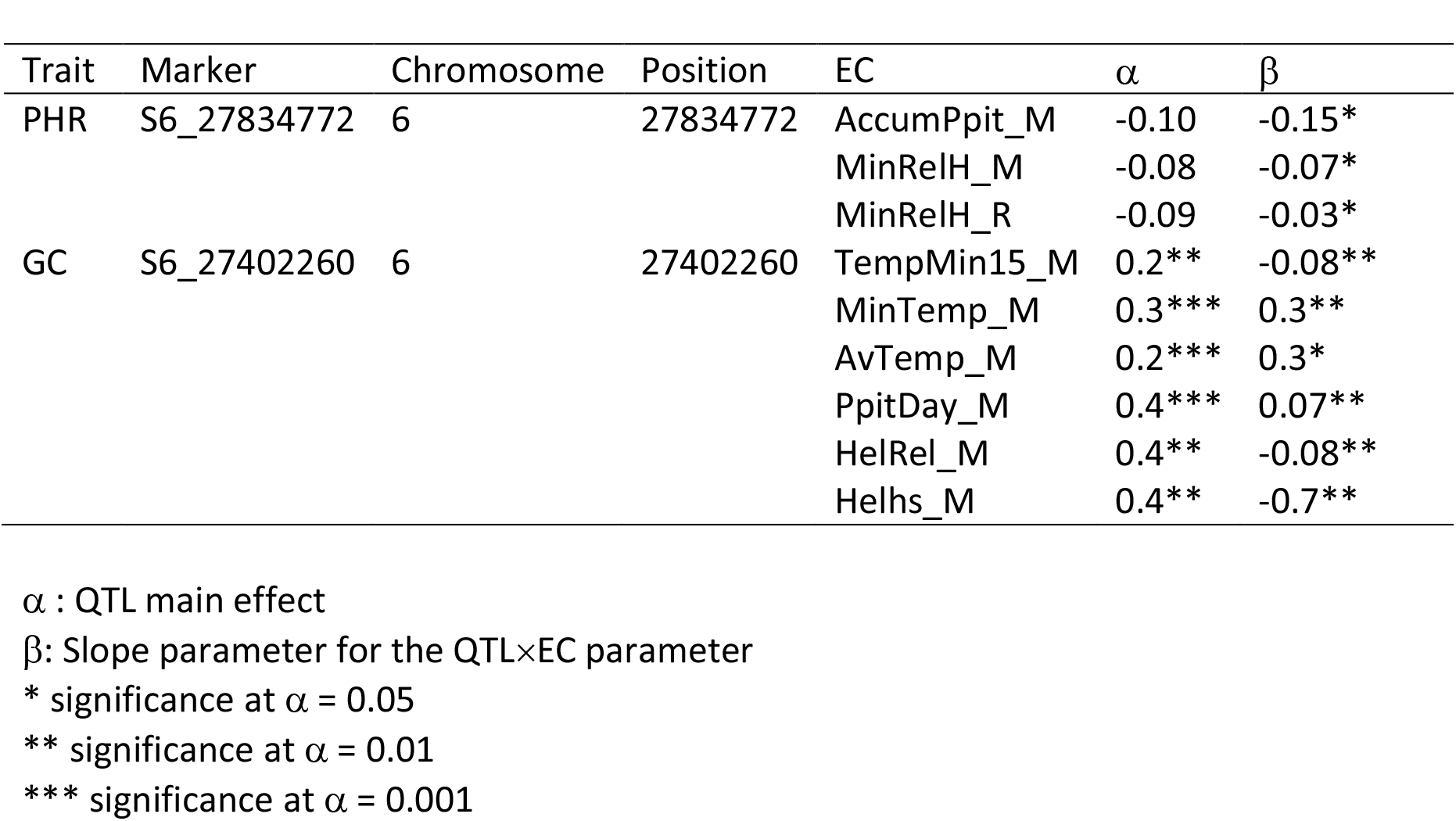
QTL responses to EC for PHR and GC in the *tropical japonica* rice population.

| Trait | Marker | Chromosome | Position | EC | $\alpha$ | $\beta$ |
| --- | --- | --- | --- | --- | --- | --- |
| PHR | S6_27834772 | 6 | 27834772 | AccumPpit_M | -0.10 | -0.15* |
|  |  |  |  | MinRelH_M | -0.08 | -0.07* |
|  |  |  |  | MinRelH_R | -0.09 | -0.03* |
| GC | S6_27402260 | 6 | 27402260 | TempMin15_M | 0.2** | -0.08** |
|  |  |  |  | MinTemp_M | 0.3*** | 0.3** |
|  |  |  |  | AvTemp_M | 0.2*** | 0.3* |
|  |  |  |  | PpitDay_M | 0.4*** | 0.07** |
|  |  |  |  | HelRel_M | 0.4** | -0.08** |
|  |  |  |  | Helhs_M | 0.4** | -0.7** |

$\alpha$ : QTL main effect
$\beta$ : Slope parameter for the QTL × EC parameter
\* significance at $\alpha = 0.05$
\*\* significance at $\alpha = 0.01$
\*\*\* significance at $\alpha = 0.001$

## Discussion

In this work we proposed to characterize and interpret G×E interaction for four traits (GY, PHR, GC and PH) in two different breeding populations of rice (*indica* and *tropical japonica*) grown in a subtropical/temperate climate. In the first part of our paper, we compare the performance of different genomic prediction models that account for genotype, environment and G×E components, to predict untested years, and we identify the most influential weather covariables for our two datasets. In the second part, we map environment-specific QTL and study the environmental variables that affect their expression, in order to interpret the QTL×E effects that account for the total G×E.

### Prediction accuracies for untested environments

Usually genomic prediction models are tested and compared using cross-validation strategies. In a multiple environment context, most studies include two basic random cross-validation schemes (Burgueño et al. 2012): CV1, which tests the performance of lines that have not been evaluated in any of the observed environments, and CV2, which tests the performance of lines that have been evaluated in some environments but not in others. These two scenarios have the disadvantage of training and validating the models with the same data, which could lead to an overestimation of the prediction accuracy the model would attain if it had been applied in an independent test dataset. Predicting new environments is a more difficult task but could represent a good validation strategy because the performance of prediction models is assessed in an independent dataset. In this work we used a cross-validation scheme for prediction in untested environments, represented by years, a component of G×E that is not easy to reproduce. This is a very relevant type of prediction for a small plant breeding program, where data from multiple locations is either limited or absent, and the need is to predict which lines are more likely to perform better in future environments. The use of EC to model the environment component explicitly has been previously shown to increase prediction accuracies for untested environments (Malosetti et al. 2016, Jarquín et al. 2017), and this situation also applies to our work.

For prediction, we compared two modeling approaches that differ in the way that multiple and correlated variables are handled: 1) a variance components approach that allows modeling the main and interaction effects of markers and ECs using covariance structures, and 2) a PLS approach that models genotype and environment effects by identifying a linear combination of all the explanatory variables, providing latent vectors that optimally predict the response variable. We found that the PLS-GW model was in all cases superior to or not different from PLS-G and reaction norm models in both datasets. Although the variance explained by the G×E component in the *indica* population was comparable in some cases to the variance explained by the genotype and/or the year main components, the proportion of variance explained jointly by the genotype, environment and G×E components, was never superior to 50% of the total variance. This could explain the lower prediction accuracies obtained in this population compared to the *japonica* population. It is possible that the EC used in this study explained only a limited proportion of the across environment interaction in the *indica* dataset, and for this reason reaction norm models, when fitting covariance matrices for the environment and marker by environment interaction, did not improve prediction accuracies in comparison to the simpler GBLUP model. In the *japonica* population, the proportion of the total variance explained by G×E was very low compared to the main genotype and environment components, which also explains why modeling a specific interaction covariance matrix did not give better results than modeling the main genotype and environment covariance matrices alone. In this population, the main environment effect was better represented by the EC, and thus, prediction accuracies, when including an EC covariance matrix (W) or the EC in the PLS model, were higher than when using a G matrix or molecular markers alone.

Besides the ability of handling numerous and correlated predictors, an additional advantage of using PLS models is that we can detect which covariates are the most explanatory in our model by looking at the model coefficients (Wold et al. 2001; Mehmood et al. 2012). Previous studies have shown the benefits of PLS for identifying the set of EC that best explain G×E (Vargas et al. 1998; Vargas et al. 1999; Crossa et al. 1999). In these studies, the G×E component of the trait was used as a response and regressed to EC only. In our case, we decided to report the results of the regression of the trait means to both EC and markers, since regressing the G×E component to EC resulted in increased MSEP with an increasing number of components, and thus a poor model fit. For GY, minimum and average temperature, and effective precipitation during flowering time showed the highest positive coefficients for *indica* rice. In regions with a temperate climate, low temperatures during flowering can affect grain yield by inducing spikelet sterility (Yoshida 1981, Alvarado 2002). The probability of occurrence of temperatures under 15°C during January (when rice usually enters the flowering stage) in Eastern Uruguay is about 20%, and would be most detrimental for *indica* varieties, which are best adapted to tropical climates. In the *tropical japonica* population, the two EC that showed the highest (negative) coefficients for GY were wind speed during flowering, and maximum temperature during grain filling. Both wind speed and high temperatures during reproduction have been proven to negatively affect GY due to pollen dehydration and consequent spikelet sterility (Marchezan and da Silva 1993; Raju et al. 2013).

For the grain quality traits, EC related to humidity, solar radiation and heliophany during grain ripening were among the most important in both datasets. The positive coefficients for solar radiation, and the negative coefficients for humidity reflect the relative effects of these variables on milling quality, as previously reported (Siebenmorgen et al. 2012; Edwards et al. 2017). Many studies have reported negative effects of high temperatures on grain chalk and percent head rice (Tashiro and Wardlaw, 1991, Lyman et al. 2013). For example, for *japonica* cultivars, temperatures higher than 26°C can cause chalky grain appearance (Chen et al. 2016), but maximum daytime temperatures higher than 33°C cause dramatic changes in the distribution of head and broken rice, and increase the proportion of chalky grain (Ambardekar et al. 2011; Lyman et al. 2013). In Eastern Uruguay, maximum temperatures during February-March, the period in which rice kernels usually develop, rarely reach 32°C. In our own dataset, the average maximum temperatures were never higher than 30°C, so it is probable that in the absence of high stress-inducing temperatures in sub-tropical rice growing areas, other variables such as humidity and solar radiation are more important, as is reflected in our results.

### QTL detection and interaction with environmental covariates

For this part of the analysis we used mixed-models to analyze QTL by EC interactions because of their flexibility, and the possibility of modeling genetic correlations between environments. We first performed an association mapping analysis for each of the four traits in each environment in both populations. In the case of PH, Rosas et al. (2017) performed a GWAS analysis on these same populations using the mean across environments and found a major effect QTL corresponding to the *sd-1* gene which was segregating in the *japonica* population, but fixed in the semi-dwarf *indica* population (Rosas et al. 2017). When we performed a single environment scan we could not find any other QTL in either population, other than a major-effect QTL corresponding to the *sd-1* gene in *japonica.*

Of the 23 QTL we found for PHR and GC in both populations, 8 were coincident with QTL reported by Quero et al. (2018) in the same populations using the mean across environments. For PHR in *indica*, we found evidence of two different genomic regions, one on chromosome 2 and another one on chromosome 6 that are affected by different weather conditions: either temperature or humidity, the two main environmental factors that affect milling quality (Cooper et al. 2008, Zhao and Fitzgerald 2013).

Two putative QTL in *tropical japonica* were co-located on chromosome 6: S6_27834772 for PHR and S6_27402260 for GC. These two QTL are in LD with qPHR.j.6.1 and qGC.j.6.2 previously found by Quero et al. (2018), and contain genes related to starch metabolism, such as *OsBEI* (LOC_Os06g51084). It is known that the expression of starch branching enzymes, like *OsBEI*, can be affected by temperature (Yamakawa et al. 2007; Sreenivasulu et al. 2014). According to our results, these two QTL showed interaction with precipitation and humidity for PHR, and with low and mean temperature, precipitation and heliophany for GC. Other researchers have shown that periods of intense solar radiation and high humidity during the ripening stage can increase the incidence of chalky grains (Wakamatsu and Tanaka 2009, Zhao et al. 2016). But these reports do not constitute proof that there is a causal relationship between the expression of these QTL and the EC, because many EC are correlated in a complex way and not all EC were observed. In temperate climates, where day and night temperatures are never as high as in the tropics, other environmental factors such as humidity and solar radiation can affect milling quality in a negative way. These findings should be confirmed by analyzing more lines in more environments to properly quantify QTL main and environment-specific effects.

In this work we used PLS, multiplicative reaction norm and mixed models to analyze our data, predict genotypic performance for yield, height and milling quality traits, and detect QTL by EC interactions. In all these analyses we assumed that the relationships between molecular markers and EC were linear, which constitutes a major limitation since interactions between genes and environmental conditions may take many different forms. A next step would be to fit statistical models with more biological realism, using models that could accommodate non-linear and more complex responses over a more extensive number of environments. Crop growth models also hold promise as a way to integrate more complex biological knowledge into the prediction process of G×E (Bustos-Korts et al. 2015, Malosetti et al. 2016). Although rather small, our two datasets allowed us to extract some broad conclusions about the nature of G×E in the Uruguayan mega-environment. Results from PLS and QTL by EC interactions indicate the effect of humidity and solar radiation on milling quality traits in temperate regions, where maximum temperatures are never as high as in the tropics. Additional research, including more environments and modeling nonlinear relationships between genes and EC, will be of particular value to better understand and predict the nature of G×E for commercially relevant traits of rice grown in temperate regions.

## Acknowledgments

The authors wish to thank the technical staff at INIA-Treinta y Tres and INIA-Las Brujas in Uruguay, and at Cornell University for their valuable support in laboratory and fieldwork. We are especially grateful to Jean-Luc Jannink and Deniz Akdemir for their valuable input and comments regarding data analysis. This study was funded by INIA’s Rice Association Mapping Project, a Ph.D. fellowship to support E. Monteverde from Monsanto’s Beachell-Borlaug International Scholarship Program, and research support from the Fulbright Commission and the Uruguayan Research and Innovation Agency (ANII).

